# Evaluation of different strategies for efficient sporulation and germination of the MICP bacterium *Lysinibacillus sphaericus* strain MB284 (ATCC 13805)

**DOI:** 10.1101/2022.09.15.508202

**Authors:** Seyed Ali Rahmaninezhad, Yaghoob A. Farnam, Caroline L. Schauer, Ahmad Raeisi Najafi, Christopher M. Sales

## Abstract

Environmental harsh conditions are one of the main challenges to the survivability of bacteria during microbially induced calcium carbonate precipitation (MICP) process. Due to the high resistivity of endospores against inhospitable conditions in comparison with vegetative cells, different sporulation methods were applied to *Lysinibacillus sphaericus* strain MB284 by changing the environmental conditions to investigate the growth of germinated cells. It was found that the sporulation yield was more when both carbon source starvation and the thermal shock process were applied to this bacterium. In addition, extending the sporulation time of cells into the minimal medium at 2 °C for a couple of weeks had a great impact on improving the sporulation yield. Comparing the growth rate of germinated endospores in natural conditions (pH 7 and 25 °C) and harsh conditions (pH 12, temperature of -10 to 60 °C, salinity up to 100 g/l) showed that endospores generated by thermal shock are able to germinate in almost every inhospitable condition except at low pH (∼3). Finally, exposing generated endospores before germination to harsh conditions (carbon source starving, high and low pH and temperature, and desiccation) for a nearly long period (to 100 days) showed that only low pH(∼3) had a negative effect on the germination process and bacterial growth curve that indicated endospore of strain MB284 can be an appropriate solution for the problem of the survivability of bioagents in MICP. These results will provide helpful information about preparing and applying endospores of *L. sphaericus* for crack healing in concrete.

**Importance:** In the bio-self-healing process, bacterial cells are responsible for the production of calcium carbonate to fill cracks in the concrete. Since cracks can happen at any time, cells must survive under harsh conditions in concrete for a long period. This study for the first time investigates different endosporulation methods to find the best well-formed endospores for microbial-inducing calcium carbonate precipitation (MICP). This study shows that the endospores of strain MB284 formed by the thermal shock can survive under inhospitable conditions including different ranges of temperatures (−4 to 60 °C), pH (3 to 14), salinity (up to 100 g/l), and starvation for about 100 days. Furthermore, the bacterial growth rate and the kinetics of calcium carbonate production by germinated endospores and vegetative cells were similar to each other that indicate endospores of strain MB284 formed by the thermal shock method developed in this study are good candidates for the MICP process.

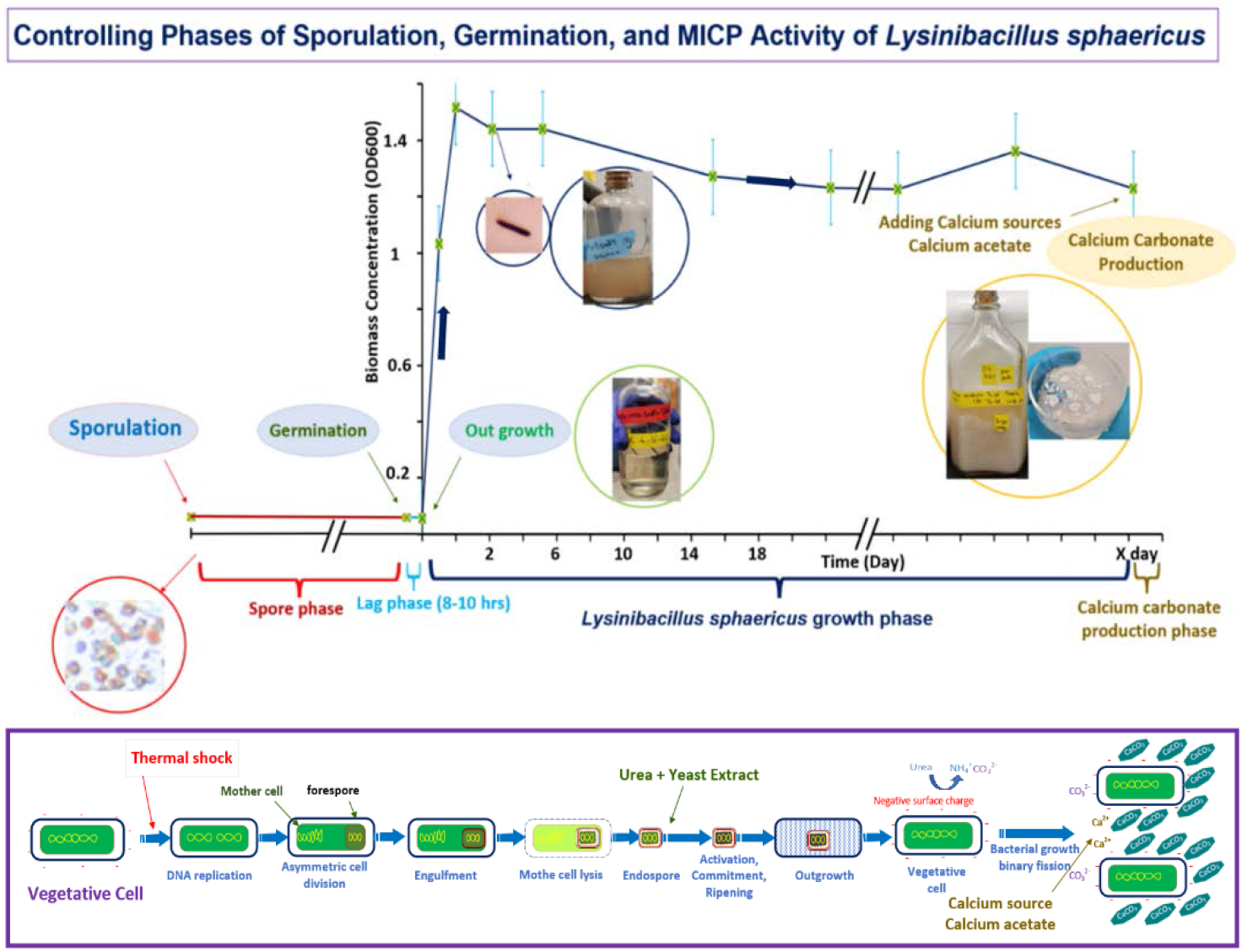

## 1. Introduction

The survivability of microorganisms used for performing microbial-induced calcium carbonate precipitation (MICP) in self-healing bio concrete application is a big challenge because, during the casting, the hardening, and service-life of concrete, microorganisms can be exposed to several environmental stresses, including high pressure, temperature, shear stress, alkaline conditions (pH ∼ 13), as well as a lack of enough oxygen, moisture, nutrient, or living space [1, 2, 3, 4]. While vegetative cells of non-spore-forming bacteria cannot survive in these inhospitable conditions, some genera of bacteria, including *Bacillus* and *Clostridium*, under such harsh conditions, can generate inactive, dormant, and multi-layered endospores. The spore-forming organisms are able to preserve their genetic content and overcome unfavorable mechanical, physical, and chemical conditions [5, 6] including extreme starvation and temperature, freezing and thawing, desiccation, high hydrostatic pressure, depletion of nutrients, and exposure to solar, ultraviolet, ϒ-radiations, organic solvents, oxidizing agents, alkaline, acids, toxic and corrosive chemicals and physical abrasion [4, 7, 8, 9, 10, 11, 12]. Although the process of sporulation is complex and largely unknown for many spore-forming species [13], studies on spores have revealed common defense mechanisms, such as coating layers, that exhibit low permeabilities and are considered the first defense lines against thermal shock [10], toxic chemicals, and enzymatic attacks [14, 10, 12]. Furthermore, there are extra layers inside the endospores such as the outer membrane, cortex, and inner membrane that enhance the adaptability of endospores to harsh conditions.

Even though spores have been proven to germinate and heal cracks in a number of studies [6, 15, 16], the long-term durability and survivability of endospores in concrete have not been fully investigated to date [17, 13]. For example, Wang et al. (2017) reported that endospores could germinate and grow in a broad range of pH and temperature, tests were performed immediately after spores were exposed to different conditions, making it unclear if spores could survive for longer periods (e.g., days to months) [16]. In addition to improving our understanding of how long spores can survive under relevant conditions in concrete, it is also important to determine under what conditions and how fast spores can germinate into active vegetative cells that can perform MICP. Studies have reported that the germination of endospores in adverse conditions can have negative effects on the viability and germination rate, lag phases, and the rate of bacterial growth and enzymatic activity [3, 2, 12]. For example, when the temperature decreased from 28 °C to 10 °C, the lag phase during the germination process increased (from 12 hr to 6 days) and the revival of ureolytic activity rate decreased (from 82.5 to 25.4 mM decomposed urea/hr) [16]. In another study, Bressuire-Isoard et al, reported that when the temperature of the environment was changed from either 45 °C or 19 °C to 37 °C, the sporulation time of *B. subtilis* decreased from 10 or 14 days to 3 days, respectively [12], suggesting that applying different environmental conditions (either naturally or manually in a laboratory) can affect the sporulation process.

Although the sporulation rate of some species has been investigated when the temperature, pH, fermentation time, rotating speed, and culture media content were manipulated [18, 19], the effect of other environmental conditions during sporulation, such as the presence of nutrients, on germination has not been tested. Therefore, in this study, the effect of applying different environmental conditions during the sporulation phase (including starvation, different culture media and nutrients, high and low temperature, pH, and bacterial concentration) on the shape of generated endospores of strain *L. sphaericus* ATCC 13805 (MB284), its sporulation and germination rates, and bacterial growth rate were investigated to determine the suitable process for endospores production.

## 2. Material and Methods

### 2.1 Culture preparation

The strain *Lysinibacillus sphaericus* ATCC 13805 (MB284) in this study was used as the bio-agent because of its great ability to produce endospores in addition to high calcium carbonate production rates [20]. For the production of the vegetative cells, strain MB284 was inoculated into the culture media containing minimal solution media (MSM) enriched by casamino acids until the biomass concentration reached OD_600_ ≈ 1. Then the bacteria was extracted from the culture medium and washed three times with phosphate buffer solution (PBS). To ensure that they are in the vegetative phase, a small portion of them were seeded into the culture medium until the OD_600_ reached around 1. For all experiments, bacteria were extracted from this culture medium. Two minimal media (MSM and EZi medium) were used in this research and their compositions have been fully described in our previous research [20].

### 2.2 Bacterial staining

The gram-staining method was based on adding crystal violet, iodine, and safranin to the bacteria that were fixed on the microscope slide. The Gram-positive bacteria stain purple while the gram-negative become red. The negative staining technique (Maevals technique) was used as the capsule staining protocol that was applied by adding congo red and crystal violet. As a result of a lack of nutrients and quorum sensing signals, vegetative cells enter the endospore-producing phase through asymmetric division during sporulation [12]. Released endospores from inside of the mother cells of the strain MB284 appear as green circles by light microscopy (Leitz Diaplan, 020-437.035) when stained by the malachite green staining technique (the Schaeffer-Fulton method). Therefore, the population and clarity of these green circles can be related to the sporulation yield and maturity of endospores.

### 2.3 Sporulation methods

The vegetative phase of strain MB284 was exposed to different environmental conditions while inoculated into the minimal media and placed into the shaker incubator and fridge at different temperatures. Acidity and alkalinity in the culture medium were applied by adding sulfuric acid and sodium hydroxide, respectively. To produce endospores based on the thermal shock process, vegetative cells were extracted and washed three times then inoculated into the MSM. The container containing vegetative cells and MSM was placed into the boiling water for 15 minutes and then immediately transferred into the ice bath for 20 minutes. Finally, they were placed in the shaker incubator for 2 days as sporulation time.

## 3. Results and Discussion

### 3.1. Physical property of strain MB284

Results from Gram-staining revealed that strain MB284 was Gram-positive (Figure 1 A), similar to other species of *L. sphaericus* [21, 2] that have thick peptidoglycan layers, where the inside of the cells are stained purple and the surrounding areas in red. Regarding capsule staining, the strain MB284 cannot create a capsule, because as Figure 1 B shows, there is no clear halo between the cells and the slide background. The absence of a capsule around the cell membrane indicates that the vegetative phase of the strain MB284 cannot produce an extracellular viscous outer layer that usually protects bacteria from desiccation and phagocytosis.

**Figure 1:**
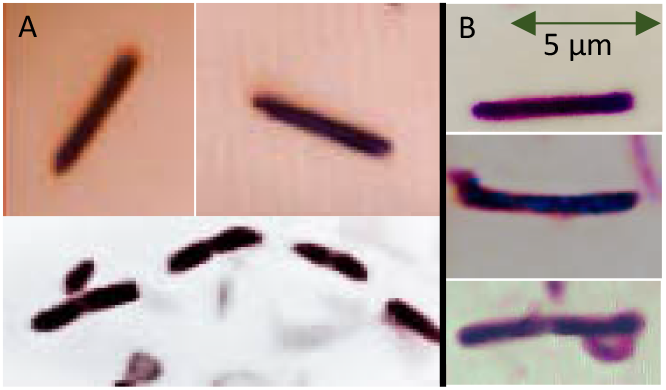
A) The strain MB284 (gram positive), B) Disability of the strain MB284 for capsul creation.

### 3.2. Impact of environmental conditions on endospore formation

Since nutrient starvation is the main and essential reason for triggering sporulation, lack of carbon sources in addition to different inhospitable conditions (high and low temperature, and pH) were applied to vegetative cells of strain MB284 to investigate the shape of generated endospores.

#### 3.2.1 Starvation

Spore-forming strains, when starved for nutrients, trigger the sporulation process to initiate dormant bacterial endospores [7]. To investigate the effect of carbon source starvation on the formation of endospores, the fresh cells of strain MB284 were transferred into the MSM at 25 °C without carbon sources. As Figure 2 A shows, some of the vegetative cells during the first day of starvation turned into the endospores (the presence of the dark green dots) indicating that carbon source starvation can initiate the sporulation process. In addition, the sporulation time was extended to one month to investigate the effect of time on endospore formation. After one month, the culture medium was diluted by phosphate buffer solution (PBS) to OD_600_≈0.1 for better observation under the electron microscope. The results showed that every single vegetative cell turned into a tiny circular endospore (Figure 2 B), which confirms strain MB284 has higher sporulation yields over time.

**Figure 2:**
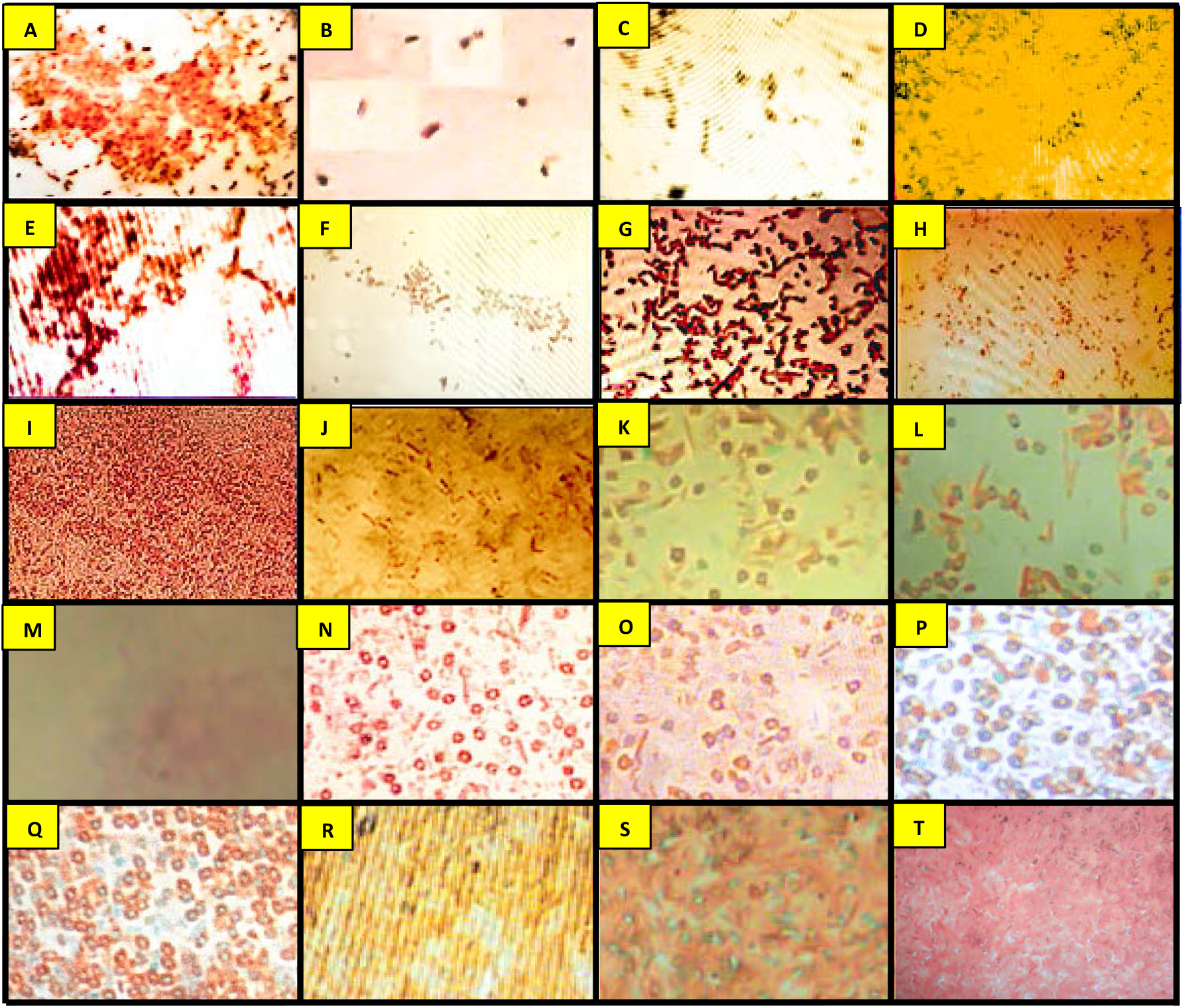
Forming spores by applying vegetative cells into different environmental conditions in the MSM; A) starvation after 1 day, B) starvation after 1 month, C) alkaline condition after 1 week, D) neutral condition after 1 week, E) storage at 45 °C for 1 week, F) storage at 35 °C for 1 week, G) storage at 2 °C for 1 week, H) -10 °C for 1 week, I) storage at 2 °C for 1 month, J) storage at -10 °C for 1 month, K) Thermal shock at neutral condition, L) Thermal shock in alkaline condition, M) Thermal shock in acidic condition, N) Thermal shock in neutral condition after one week, O) Thermal shock at alkaline condition after one week, P) Thermal shock in neutral condition after one month and stored at – 2 °C, then washed 5 times by autoclaved ultrapure water Q) Thermal shock in alkaline condition after one month and stored at – 2 °C, then washed 5 times by autoclaved ultrapure water, R) Thermal shock in neutral condition without inoculation, S) Thermal shock in neutral condition inoculation in the MSM, T) Thermal shock in neutral condition inoculation in the MSM enriched by casamino acids.

#### 3.2.2 pH

Since the bacterial activity and morphology of cell membranes are affected by different hydrogen ions concentrations [22], exposing bacteria to different pH may impact the shape of endospores. Therefore, acidic (pH=3), alkaline (pH=12), and neutral conditions (pH=7) were applied to the vegetative cells inoculated into the MSM at 25 °C to investigate the effect of pH in addition to starvation on the shape of endospores and sporulation yield. Based on the results, no vegetative cells and endospores were observed in acidic conditions after one week of inoculation, revealing the sensitivity of both cells and spores of strain MB284 to lower pHs (Picture was not shown). However, comparing Figures 2 C & D it was observed that partial sporulation, at near similar rates, happened in both neutral and alkaline conditions, indicating that sporulation rates are not significantly different at pH 7 or 12.

#### 3.2.3 Temperature

Although it is documented that the ideal growth temperate for *L. sphaericus* species is 30 °C [23], it is unknown if the temperature can have an impact on the sporulation of strain MB284. Therefore, cells of strain MB284 were starved and stored in the MSM at different environmental temperatures (−10, 2, 35, and 45 °C) to investigate the effect of temperature on the shape of the resultant endospores. Results showed that after one week of inoculation when the temperature was 45 °C (Figure 2 E) more endospores were generated than at 35 °C (Figure 2 F). And, even though the lower temperature of 2 °C (Figure 2 G) also led to higher sporulation than at 35 °C, decreasing the temperature down to -10 °C (Figure 1 H) had the worst sporulation yield. Furthermore, extending the sporulation time to one month revealed that the sporulation yield increased at both 2 and -10 °C (Figure 2 I & J). We hypothesize that the rate of sporulation at 2 °C was more than -10 °C due to the slower metabolism of vegetative cells under freezing conditions (−10 °C).

Since high and low (close to zero degree centigrade) temperatures caused strain MB284 to generate more endospores, a cycle of high and low temperature was applied to vegetative cells of strain 284 inoculated into the MSM to investigate the effect of thermal shock on generating endospores. The result showed that thermal shock caused vegetative cells to generate fully developed, well-structured, and circular endospores (Figure 2 K). The separated and clearly detectable endospores indicate that they were thoroughly released from the mother cells to the culture medium. A similar method to thermal shock was previously applied to *L. sphaericus* cells by Intarasoontron et al. [4], but the shape of generated endospores in that study was not presented and compared with other sporulation methods.

During the inoculation period of the thermal shock process, alkaline (pH=12) and acidic (pH=3) conditions were applied to the MSM to investigate the effect of pH in addition to starvation and on the shape of endospores in the thermal shock method. The effects of pH were similar to section 3.2.2 when sporulation occurred in environmental temperature (25 °C). The shape and number of endospores in the alkaline condition (Figure 2 L) were similar to the neutral condition, however, no endospores were formed in the acidic condition (Figure 2 M). The sporulation period was extended to one week (instead of two days) in both neutral and alkaline conditions while the samples were stored at 2 °C to investigate the effect of sporulation time on the shape of generated endospores. It was observed that the number of endospores increased in both neutral (Figure 2 N) and alkaline (Figure 2 O) conditions indicating the effective role of time in generating endospores that are well aligned with the result of section 3.2.1. Regarding the enhancing effect of sporulation time on generating endospores, samples in both neutral (Figure 2 P) and alkaline (Figure 2 Q) conditions were kept for one month at 2 °C and finally were washed five times with autoclaved ultrapure water. Based on the result, washing by water isolated endospores from remaining dead cells, extracellular polymeric substances (EPS), and soluble microbial products (SMP). The well-shaped, packed, and highly concentrated endospores generated in both neutral and alkaline conditions proved that thermal shock after one month of inoculation at 2 °C and finally washing 5 times by autoclaved ultrapure water is an appropriate method for isolating dense concentrations of endospores.

To investigate the influence of the inoculation process in the thermal shock method on the endospores formation, a sample was taken after the ice bath step and before inoculating in the MSM. The result showed that no endospores were observed in this step (Figure 2 R), which illustrates that inoculation is essential for strain MB284 to generate multilayered endospores.

In our previous study [20], we found that strain MB284 requires some unknown amino acids, such as those in a casamino acid mixture, to grow. To investigate the effect of starvation on making endospores, 20 g/l of casamino acids as a carbon source was enriched in the MSM during the sporulation time in the thermal shock method. A comparison between the number of generated endospores inoculated into the MSM (Figure 2 S) with that of MSM enriched by casamino acids (Figure 2 T) after three weeks showed that adding casamino acids during inoculation had an adverse effect on the sporulation process. The presence of carbon sources facilitated germination, thus reducing the number of generated endospores. While a medium containing peptone and beef extract has been used in some attempts for making endospores [3], comparing Figures 2 S and T revealed that starvation had a positive role in generating endospores for the strain MB284.

### 3.3 Effect of different environmental conditions on endospore germination

Dormant endospores are able to sense their surrounding environment, and whenever the environment becomes favorable and suitable for their growth, they break dormancy and initiate germination and outgrowth processes to regain the physiology and metabolic functions of vegetative cells [7, 9, 10, 12]. The length of the lag phase during the germination process depends on the type of species and environmental condition and can vary from a couple of minutes to more than a couple of days [8]. Endospores in concrete during their dormancy may experience several environmental harsh conditions (such as high and low temperature, pH, and desiccation) that affect the germination process, outgrowth length, and MICP activities. Therefore, the effect of different environmental conditions during the dormancy period and germination time on the bacterial growth curve was investigated in this section.

#### 3.3.1. Culture media

In our previous research [20], we noticed that strain MB284 is auxotrophic and cannot grow in defined minimal media without casamino acids or casein or other nutrient-rich sources, such as peptone, beef, yeast, and meat extract. Among these nutrient-rich sources, we found that yeast extract was the optimum undefined culture medium for strain MB284 to grow and produce calcium carbonate. Therefore, in this experiment, the germination of endospores was investigated in three different culture media: MSM with either casamino acids or casein, or yeast extract in DI water that all contained 20 g/l of urea.

The results from this experiment showed that there is no difference in the bacterial growth curve when endospores were germinated in either 10 g/l of casamino acids or yeast extract (Figure 3 A). However, the bacterial growth rate in the exponential phase was lower in casein (0.66 OD_600_/d) rather than that of both casamino acids (1.36 OD_600_/d) and yeast extract (1.42 OD_600_/d), while the maximum biomass concentration (based on OD_600_ as an indication of biomass concentration) was higher in casein (2.09) rather than that of casamino acids (1.62) and yeast extract (1.66). The bacterial growth curves we observed for endospores inoculated into the casamino acids, yeast extract, and casein were highly similar to the vegetative bacterial growth curve we found in our previous study [20] indicating the germinated endospores grew similar to the culture started with vegetative cells. We hypothesize that the greater growth rates with casamino acids and yeast extract than casein is due to their higher solubility in water, which makes the bioavailability of nutrients and energy sources in media containing casamino acids or yeast extract initially higher leading to faster growth. Although casamino acids and yeast extract have faster initial growth rates, the growth potential of strain MB284 with casein compared to casamino acids, at the same concentration (10 g/l), is higher.

**Figure 3:**
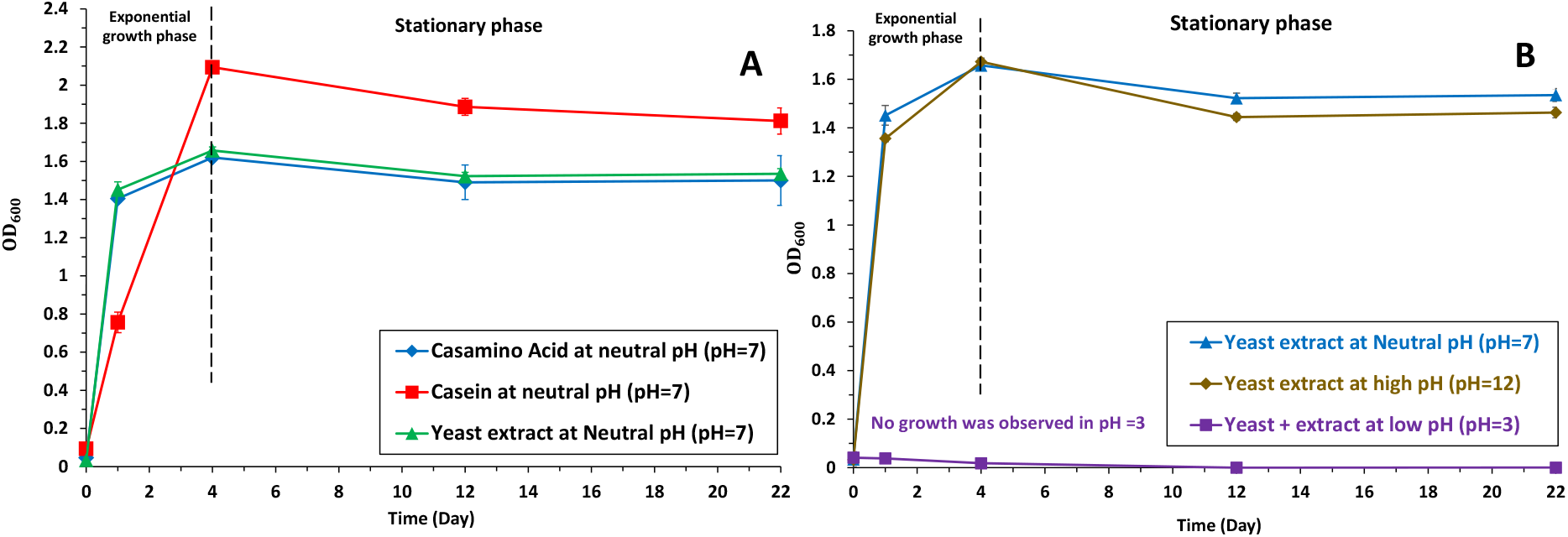
Endospore’s germination in the MSM enriched either by Casamino acids or Casein, or Yeast extract in DI water at low pH (pH=3), neutral (pH=7), and high pH (pH=12)

#### 3.3.2. pH

The germination of strain MB284 in yeast extract in different pH conditions indicated that endospores cannot germinate and outgrowth in acidic conditions (pH=3) after 22 days (Figure 3 B). Thus, the lower pH is not a suitable condition neither for sporulation (based on the results of section 3.2) nor germination of strain MB284. However, since the micro-environment in concrete is either neutral or alkaline conditions, the disability of strain MB284 to outgrow in the acidic condition is not considered a challenge for the bio self-healing process.

Although, Wang et al. [16] reported that increasing pH extended the lag phase of the germination of *B. sphaericus LMG 22257* to 40 hours, comparing the growth curve of strain MB284 inoculated in yeast extract in neutral (pH=7) and alkaline (pH=12) conditions indicated that high pH condition did not extend the lag phase of germination process (Figure 3). In addition, the exponential growth rate in the alkaline condition was 1.32 OD_600_/d and the maximum biomass concentration based on OD_600_ was 1.67 that are very close to the neutral condition values mentioned in section 3.3.1.

#### 3.3.3. Nutrients

Two minimal media (the MSM and EZi medium) with different ion contents were prepared and enriched by casamino acids containing carbon source and required amino acid for the growth of strain MB284. The endospores were inoculated into these two media to investigate the impact of the types and concentration of ions on the germination process. As Figure 4 illustrates, the required time for germination of endospores in the EZI medium was considerably more than the MSM. In addition, the bacterial growth rate and the maximum biomass concentration were greater in the MSM compared to the EZi medium. Based on our previous research [20], the growth curves of vegetative cells inoculated into the MSM and EZi media were similar, therefore the current significant difference between the growth curves of bacteria belonging to the MSM and the EZi media was due to the endospore’s nutrient requirement during the germination process. Endospores uptake ions during the outgrowth step, therefore, since the diversity and concentration of salts in the MSM (especially the presence of Mn^2+^, Mg^2+^, Zn^2+^, and Ca^2+^ that has a positive role in germination [12]) were more than the EZi medium, it was found that the germination of strain MB284, highly depends on the concentration of ions.

**Figure 4:**
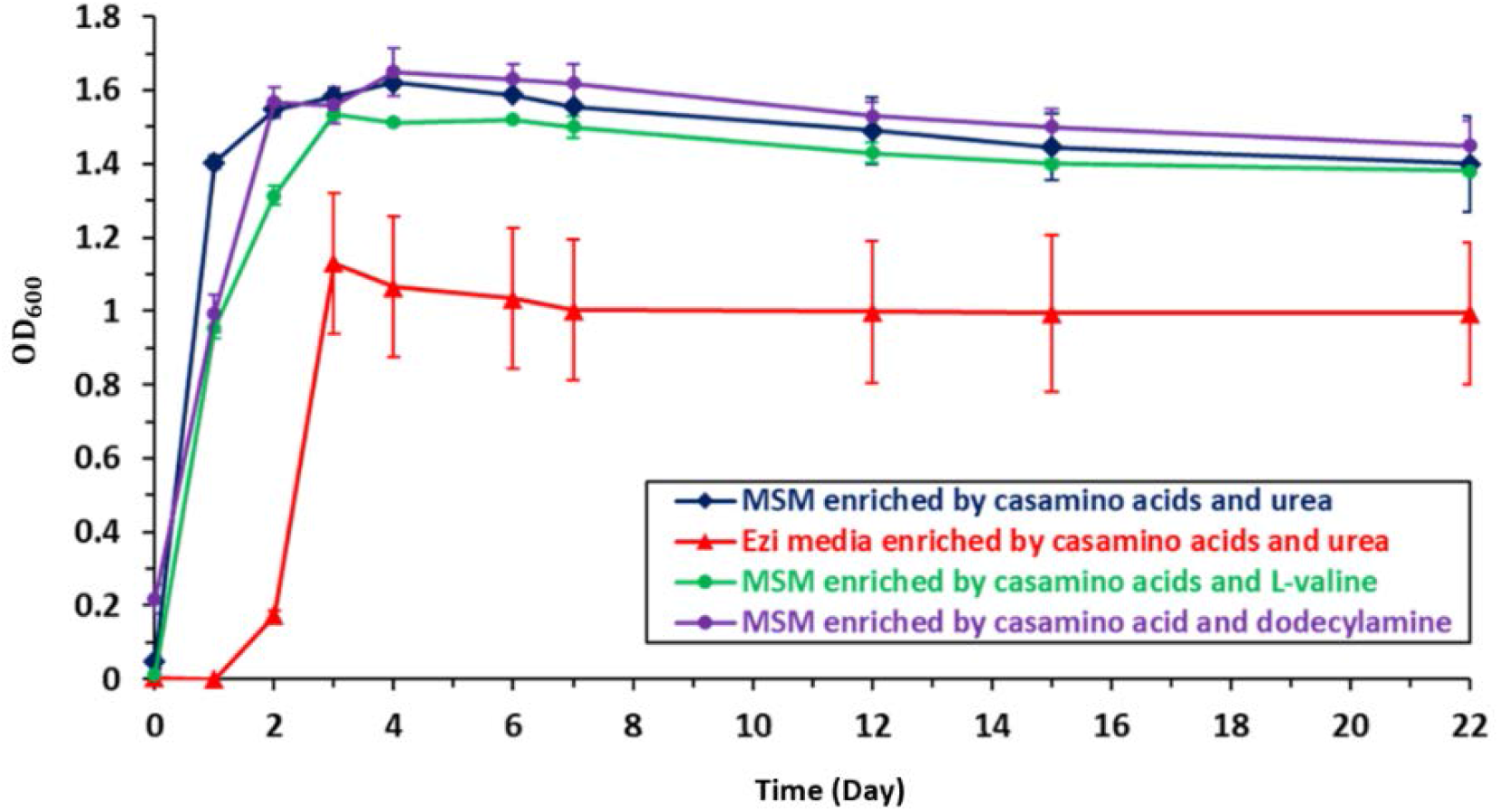
The growth curve regarding the germination of endospores in the MSM and EZi media enriched by casamino acids and also in the MSM enriched by casamino acid and either L-valine or dodecylamine.

#### 3.3.4. Germination aiding agents

In addition to adding nutrients, non-nutrient agents such as dodecylamine, CaDPA, L-valine, L-alanine, L-proline, L-asparagine, and surfactants can trigger endospore germination [12, 9]. Although the lag phase for germination of the strain MB284 was not long in the absence of aiding agents, the impact of L-valine and dodecylamine on the growth curve was investigated in this study (Figure 4). Comparing the growth curve of strain MB284 inoculated into MSM containing casamino acids and either L-valine or dodecylamine revealed that these two aiding agents do not decrease the lag phase duration or increase the growth rate and maximum biomass concentration.

#### 3.3.5. Salt concentration

The salinity of pore solutions in concrete varies depending on the salt content of the water and the sand which are used in its production, which can be as high as ∼ 90 g/l when seawater and sea-sand are used [24]. Although growth was observed to be higher in MSM, which has a higher concentration of ions than the EZi medium (Figure 4), it is unclear how halotolerant strain MB284 is [25]. To investigate the effect of different salt concentrations (up to 100 g/l) on the growth rate of germinated cells of strain MB284, endospores were inoculated into the MSM enriched by casamino acids, urea, and different salt concentrations, and the biomass concentration was measured at different time intervals (1^st^, 2^nd^, 6^th^, 12^th^, and 27^th^ day). As Figure 5 A shows, after one day of inoculation, the higher concentration of salt in the culture medium had a negative impact on the bacterial growth of cultures of strain MB284 when inoculated with spores, with no growth observed with salt concentrations greater than 30 g/l. However, it was observed that beyond 1 day, cultures of strain MB284 were eventually able to grow in higher salt concentrations with an extended inoculation period. For example, germinated spores of strain MB284 had noticeable growth after 27 days when the salt concentration of the medium was around 90 g/l (Figure 1A-2).

**Figure 5:**
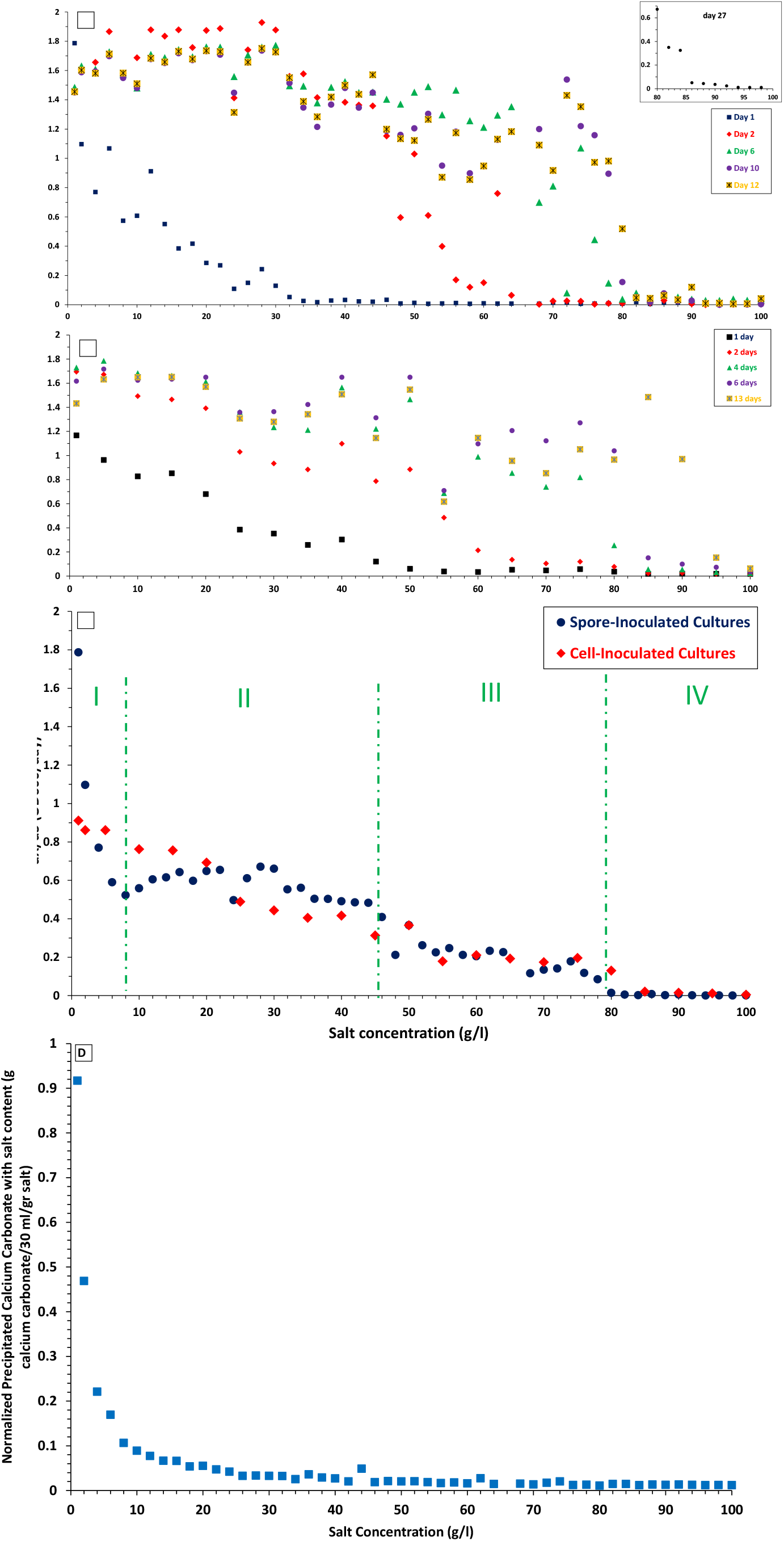
Investigation of the effect of salt concentration on the biomass of A) germinated endospores and B) vegetative cells. Growth kinetics of C) germinated endospores and D) vegetative cells (in the formula C and S represent the biomass concentration in OD_600_ and salt concentration in g/l. The produced calcium carbonate normalized with added salt.

To compare the performance of the growth of germinated endospores in different salt concentrations with cultures inoculated with vegetative cells, the vegetative phase of strain MB284 was inoculated into the MSM enriched by casamino acids in different salt concentrations (up to 100 g/l) and biomass concentrations were measured over different time intervals (1^st^, 2^nd^, 4^th^, 6^th^, and 13^th^ day) (Figure 5 B). The comparison between Figures 5 A and B reveals that even though higher salt concentrations had a negative effect on both cultures of strain MB284 when inoculated by either vegetative cells or germinated endospores, vegetative cells were able to grow faster than germinated endospores in higher salt concentrations. For example, after one day of inoculation, the only condition where cultures of strain MB284 when inoculated with vegetative cells were not able to grow was with salt concentrations greater than 50 g/l. Similar to cultures of germinated spores, cultures inoculated with vegetative eventually demonstrated growth after 13 days, even with 90 g/l of salt.

The lower growth rates of cultures inoculated with endospores in low salt concentrations compared to those inoculated with vegetative cells (Figure C) is hypothesized to be due to the time required by the spore to germinate. 20 g/l of calcium acetate was added to culture media with different salt concentrations to investigate the effect of salt on calcium carbonate production. Since the concentration of germinated endospores after 27 days of inoculation was nearly similar, the produced calcium carbonate was normalized with added salt (Figure 5 D). An increase in the salt concentration decreased the production of calcium carbonate that is because of the presence of cations (Na^+^) in addition to Ca^2+^ that interact with the surface of bacteria and decrease the available nucleation sites for the production of calcium carbonate on the bacterial membrane cell.

#### 3.3.6. Effect of storage environmental conditions on germination of endospores

During the casting, hardening, and service life of concrete, endospores experience different severe conditions that may affect the germination process. To investigate the effect of these harsh environmental conditions on the germination and growth curve, the endospores of strain MB284 were exposed to different temperature and pH conditions prior to germination. Specifically, endospores were kept in the acidic (pH=3), neutral (pH=7), and alkaline (pH=12) conditions in the MSM culture medium for 7 and 21 days. Then endospores were extracted and washed 3 times by PBS, inoculated into the MSM enriched by 20 g/l of urea and yeast extract. The results showed that keeping endospores in the alkaline condition both for 7 and 21 days had no significant effect on the bacterial growth rate. While endospores lost their germination ability when they were stored in the acidic condition (Figure 6 A).

For evaluation of the effect of temperature, freeze-thaw cycles are one of the main reasons for creating fractures during concrete service life. Thus, it is very common that endospores to experience high and low temperatures while there are in the concrete. Therefore, endospores were inoculated into the MSM enriched with 20 g/l of urea and stored at 10 °C and 50 °C to investigate the effect of keeping endospores at different temperatures on the germination rate. After 21 days, 10 g/l of casamino acids were enriched into the medium to trigger the germination at 30 °C. The results indicated that storing endospores in low and high temperatures had no negative effects on the germination process and growth curve (Figure 6 A).

In addition to being exposed to extreme temperatures and pHs, it is also possible that the endospores of the bioagent could be poorly distributed throughout the concrete, creating microenvironments within the concrete where fractures may occur with varying concentrations of endospores. Therefore, the effect of endospores concentration on germination rate was also investigated in this study. Specifically, two different concentrations of endospores (0.053 and 0.35 OD_600_) were stored for 15 days in the MSM, then enriched with casamino acids for triggering the germination phase. As Figure 6 B shows, the bacterial growth rate for the high endospores’ concentration (1.13 OD_600_/day) was a little higher than that of the fewer endospores’ concentration (1.06 OD_600_/day), however, two culture media finally possessed the same biomass concentration after a couple of days of germination.

Another factor that may affect germination that was investigated in this study is the age of the endospores. The endospores were stored in different bottles containing the MSM and germinated in different periods. Although endospores are required to effectively remain alive for even more than 100 years, regarding the laboratory limitation, the maximum storage time in this experiment was 100 days. The results showed that the germination rate was quite similar when endospores were germinated in different time intervals and able to germinate at least up to 100 days of sporulation. Lastly, since the humidity of concrete may change and decrease over time, it is possible that the endospores may experience desiccation. Therefore, to investigate the effect of lacking water on germination, endospores were dried in the oven at 35 °C and stored for 70 days at 40, 30, and 2 °C, then were inoculated into the MSM enriched by 20 g/l of casamino acids and urea. As Figure 6 D illustrates, storing dried endospores at 2 °C decreased the growth rate in comparison with storing at 30 and 40 °C.

In conclusion, this paper demonstrates that applying carbon starvation and thermal shock to vegetative cells of strain MB284 can effectively lead to the generation of endospores that can tolerate different harsh environmental conditions for a long period (up to 100 days). The paper also revealed that exposing these endospores to carbon and nutrient sources, such as yeast extract, leads to their rapid germination and urea enhances the amount of produced calcium carbonate. The findings from this paper provide methodologies for generating endospores of strain MB284 and for germinating those endospores for MICP activity so that they can be used in bio-self healing concrete applications. Since all of the experiments conducted in this study were in-vitro assays, the next step in advancing the application of endospores for self-healing would be to examine the survivability and germination process of endospores of strain MB284 in concrete.

## Acknowledgment

This material is based upon work supported by the National Science Foundation (NSF) under Grant No 2029555 so we would like to appreciate NSF for its support. Also, we would like to appreciate Asa Lewis for revising this manuscript.

